# *Toxoplasma gondii* ROP18 Inhibits Human Glioblastoma Cell Apoptosis through Mitochondrial Pathway by Targeting Host Cell P2X1

**DOI:** 10.1101/383638

**Authors:** Li-Juan Zhou, Min Chen, Cheng He, Jing Xia, Cynthia Y. He, Sheng-Qun Deng, Hong-Juan Peng

## Abstract

It is known that *Toxoplasma gondii* infection both initiates and inhibits host cell apoptosis through different proapoptotic signaling cascades, but the parasitic factors involved in these processes remain unclear. *T. gondii* virulence factor ROP18 has been reported to regulate host cell apoptosis, but the results of this regulation are few reported and contradictory. In this study, we found that immune or neuro cells infected by any one of the *T. gondii* strains (RH-type I, ME49-type II, and VEG-type III) showed a significantly lower apoptosis index than their uninfected controls when apoptosis was induced by staurosporine (STS). We further found that ROP18 of RH strain inhibited ATP induced apoptosis in human glioblastoma cells (SF268) with endogenous expression of human proapoptotic protein purinergic receptor 1 (P2X1), but had no effects on the immune cells of RAW264.7 and THP-1 without detectable P2X1 expression, which may indicate that ROP18’s inhibition of host cell apoptosis is related to P2X1. Interestingly, we further identified that ROP18 (RH strain) interacted with P2X1, and over-expression of ROP18 in COS-7 cells inhibited the cell apoptosis mediated by P2X1. We also found that ROP18 of RH strain inhibited P2X1-mediated Ca^2+^ influx, translocation of cytochrome C from mitochondria to cytoplasm, and 1 ATP-triggered caspases activation. Collectively, these findings supported that ROP18 inhibited the host cell apoptosis through the intrinsic mitochondria pathway by targeting host cell P2X1, thereby suggesting a sensor role of the host proapoptotic protein P2X1 in this process

**Author summary:** The obligate intracellular protozoan *Toxoplasma gondii* has been shown to modulate cell apoptosis through different apoptotic pathways. However, the consequences are various and even contradictory, and the parasite effectors and the precise biological mechanisms remain unclear. Herein we showed that *T. gondii* of type I, II, and III strains could inhibit the apoptosis of neuro cells and immune cells. *Toxoplasma gondii* ROP18 (RH strain) inhibited apoptosis of human glioblastoma cell SF268 by targeting C terminal of host cell P2X1 protein, but not through proteasome-dependent degradation of P2X1.

## Introduction

*Toxoplasma gondii,* an obligate intracellular protozoan, infects the nucleated cells of all warm-blooded animals including humans [1]. *T. gondii* infection shows no or mild symptoms in immune competent hosts, but the symptoms may be severe in immunocompromised patients presenting with all types of toxoplasmosis, or in case of primary infection during pregnancy which may be vertically transmitted to fetus leading to fetus deformity, abortion or newborn’s toxoplasmosis [2]. *T. gondii* strains are categorized into highly virulent type I, and non-virulent types II and III based on their acute virulence in mouse model [3]. The lethal dose (LD) of type I strain (RH) is one parasite, while the median lethal dose (LD50) of nonvirulent types II and Π strains (PLK and CEP) is more than 100 parasites [4].

Apoptosis, known as type I programmed cell death, is a biological event induced by various physiological or pathological stimuli and then processed by activation of a series of protein cleavage enzymes known as caspases [5]. Staurosporine (STS) is a protein kinase C (PKC) inhibitor, which has been used in the induction of cell apoptosis dependent on p38 MAPK pathway [6]. As an apoptosis inducer, STS induces cell apoptosis via elevating the cytosolic ATP level [7]. Adenosine triphosphate (ATP) is regarded as an energy storage in vivo and neurotransmitter in the nervous system, and extracellular ATP can induce SH-SY5Y cells apoptosis through decreasing the expression of anti-apoptotic protein Bcl2 and increasing the expression of proapoptotic protein Bax [8].

*T. gondii* is reported to both promote and inhibit host cell apoptosis, these opposing effects might involve complicated factors that modulate the exquisitely balanced interaction between the parasite and the pro-and anti-apoptotic signals of the host, such as the cell type, the virulence of *T. gondii* and so on [9]. For example, tachyzoites of the RH strain promotes the apoptosis of mouse neural stem cells [10], but inhibits the apoptosis of human leukemic cells THP-1 and Jurkat cells [11,12]. Apoptosis of trophoblast cells can be induced by ME49 infection but inhibited by RH infection [13]. Meanwhile, the major virulence factor ROP18 of *T. gondii* was also reported to induce the apoptosis of mouse neural cells N2a [14], and to suppress apoptosis in human epithelial cell 293T [15].

Human purinergic receptor 1 (P2X1) is an ATP-gated ion channel formed with trimeric assembly of the subunits with two transmembrane regions, the intracellular amine and carboxyl termini and a large extracellular ligand binding loop [16]. P2X1 receptors have a conserved intracellular serine/threonine PKC (protein kinase C) site, which functions in its potentiation through phosphorylation of an interacting regulatory protein diacylglycerol (DAG) generated by G-protein-coupled receptors [17,18]. Immunohistochemistry and functional studies have shown that P2X1 is expressed predominantly on smooth muscle cells, blood cells and neurons cells [19–21]. P2X1 activation results in Na^+^ and Ca^2+^ influx and K^+^ efflux across the cell membrane, which leads to depolarization of the plasma membrane and an increase of the intracellular Na^+^ and Ca^2+^ concentration [21] P2X1 abnormality is implicated in many diseases. For instance, the accumulation of alpha-synuclein in susceptible neurons through P2X1 mediated lysosomal dysfunction may account for Parkinson’s disease [19]. P2X1 is activated in the motoneurons in nerve injury [22]. Male infertility occurs in P2X1 knockout mice [23]. The cell apoptosis rate is increased in cells over-expressing P2X1,which indicates P2X1 is a proappoptotic protein [24]. In this study, we found *T. gondii* infection inhibited host cell apoptosis induced by STS regardless of strain virulence, *T. gondii* ROP18 targeted host cell P2X1 and inhibited ATP stimulated cell apoptosis through mitochondria pathway.

## Results

### 1-Infection of the three types of *T. gondii* strains (RH, ME49, VEG) inhibits the host cell apoptosis induced by staurosporine

Human glioblastoma cells (SF268) were infected with *T. gondii* strains RH, ME49 or VEG for 2hrs and 22hrs respectively. After infection, the control and infected groups were then treated by staurosporine (STS) for 4hrs and 6hrs respectively to induce apoptosis. Our flow cytometry (FCM) results indicated that the apoptosis indices in SF268 cells infected by different strains of *T. gondii* were significantly lower than which in the uninfected control, at both 6hrs and 28hrs post infection (PI) (Figure 1A&C). DNA fragmentation was not detected in both infected or uninfected SF268 cells after induction with STS (data not shown).

**Figure 1.**
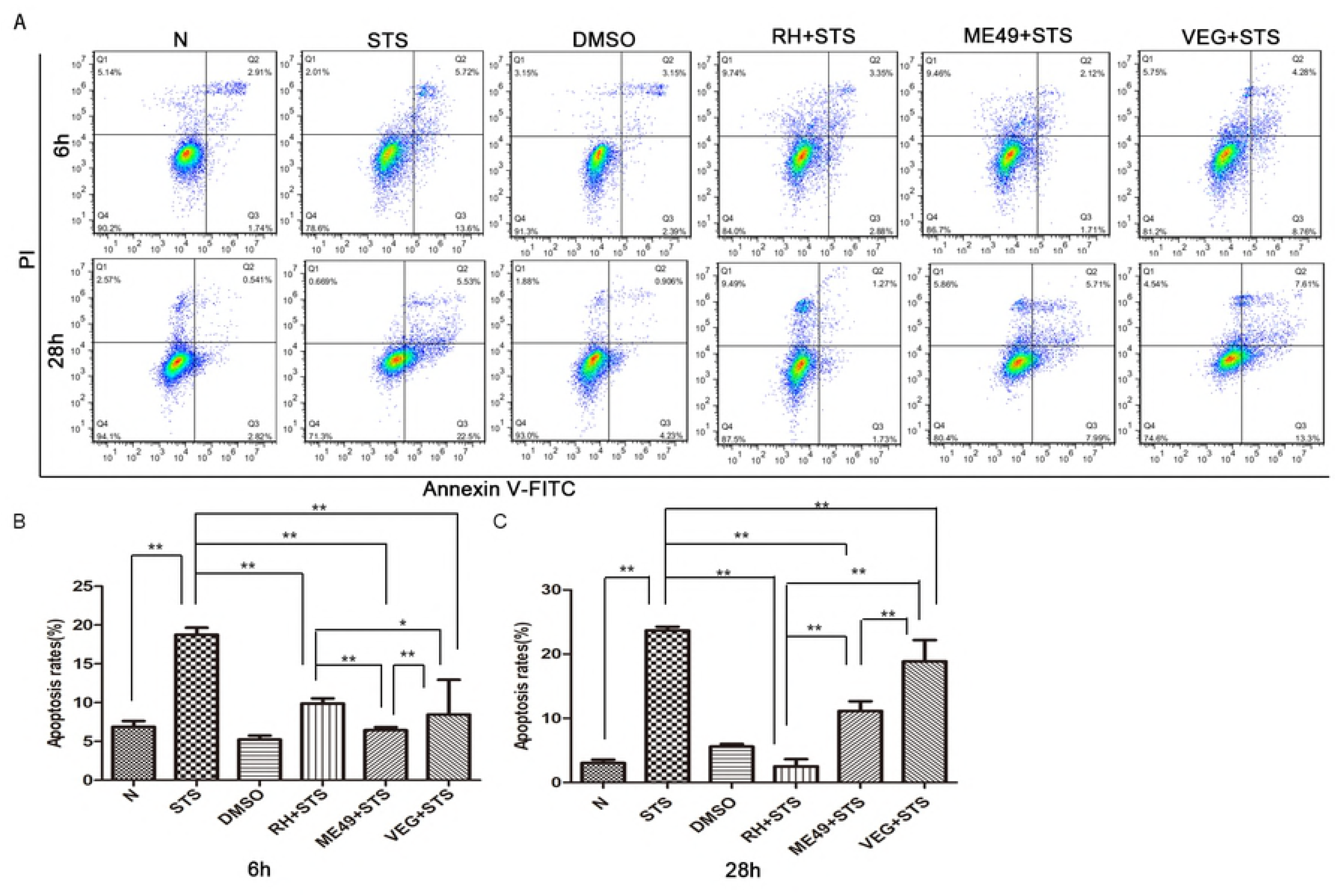
RH, ME49 and VEG infection inhibited SF268 cell apoptosis induced by STS at 6h and 28h post infection. SF268 cells were infected with RH, ME49 and VEG strains, respectively (MOI = 3) for 2h or 22h followed by treating with staurosporine (STS) for 4 h or 6h. The cells were collected for apoptosis detection by flow cytometry (FCM). A: FCM detection of apoptotic cells at 6h and 28h post infection. Apoptotic cells included the early apoptotic cells (Annexin V^+^/ PI^−^) and the late apoptotic cells (Annexin V+/ PI+). **B&C:** The quantitative data converted from FCM, at 6h and 28h post infection, respectively. Values were expressed as mean ± SD in each group, and the experiments were repeated three times for one-way ANOVA statistical analysis (**P < 0.01). The results showed that *T. gondii* infection significantly inhibited SP268 cell apoptosis induced by STS regardless of strain virulence.

Meanwhile, the other two types of cells, human monocyte/macrophage cells (THP-1) and murine macrophage cells (RAW264.7) were also used in our experiment to show the host cell apoptosis modulated by different types of *T. gondii* infection after STS induction for apoptosis. The same results were observed as in the SF268 cells (Figure S1 A&B and Figure S2 A&B). Detection of DNA fragmentation in THP-1 cells and RAW264.7 cells also showed that the relative abundance of small DNA fragments was less evident in RH, ME49 or VEG infected groups compared to their uninfected controls after induction with STS (Figure S1 C, Figure S2 C).

These results suggested that STS-induced cell apoptosis in human SF268, THP-1 and mouse RAW264.7 cells were significantly inhibited by *T. gondii* infection regardless of strain types, at both 6hrs and 28hrs PI.

### 2-*Tg*ROP18 is a regulator for suppression of ATP induced apoptosis in SF268 cells

Based on previous literature review, we found that *Tg*ROP18 of RH strain modulated host cell apoptosis, but the modulation results were related to the types of cells they infected. To find out whether the apoptosis of SF268, THP-1 and RAW264.7 cells were influenced by *Tg*ROP18, the cells of SF268, THP-1 and RAW264.7 were infected with RA-Δ*ropl8* or RH wild type tachyzoites at multiplicity of infection (MOI) of 13 (MOI=13) for 12hrs followed by ATP induction for 12hrs to induce apoptosis. The normal and ATP treated cells were used as negative and positive control, respectively. Our flow cytometry (FCM) results showed that the apoptosis rate of RH infected SF268 cells was (4.8±0.74) %, which was significantly lower (P < 0.01) than which of (11. 8±1.34) % in the RH-Δ*ropl8* infected SF268 group (Figure2A&B). While this phenomenon was not observed in THP-1 and RAW264.7 cells, both the RH and RH-Δ*ropl8* infected groups showed lower apoptosis rate when compared to the ATP treated group, but no significant difference was observed between the RH and RH-Δ*ropl8* infected groups (Figure S3 A&B). These results indicated that *Tg*ROP18 played no observable role in modulating the apoptosis of THP-1 and RAW264.7 cells, while apparently inhibited the apoptosis of SF268 cells.

**Figure 2.**
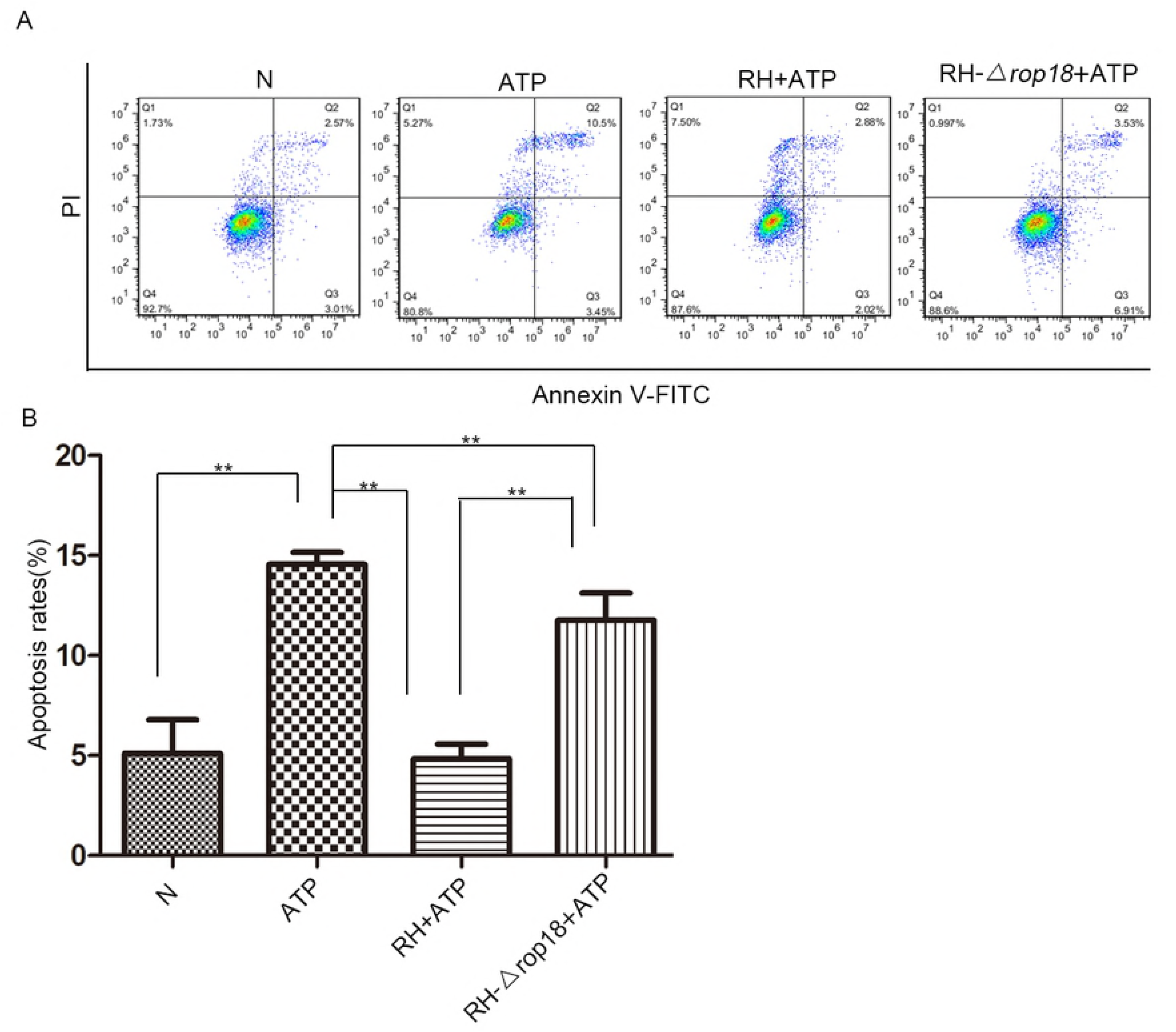
*T. gondii* major virulence factor ROP18 inhibited SF268 cell apoptosis induced by ATP. The SF268 cells were infected with RH or RH-Δ*rop18* tachyzoites (MOI=13) or left uninfected for normal control (N) or positive control (ATP treatment). At 12 h post infection, 1 mg/ml ATP was added to the cells for additional 12 h except the normal control group. A: The apoptosis of the SF268 cells from each group was detected by flow cytometry after Annexin V-FITC/PI staining. B: The percentages of apoptotic cells were separately determined for each group of cells. Values were expressed as mean ± SD in each cell group, and the experiments were repeated three times for one-way ANOVA (**P < 0.01). The SF268 cells infected by RH strain showed a significantly lower apoptosis rate compared to the RH-Δ*ropl8* infection group, suggesting that *Tg*ROP18 inhibited SF268 apoptosis significantly.

### 3-*Tg*ROP18 targeted host cell protein P2X1

To understand the molecular mechanism of *Tg*ROP18 inhibiting cell apoptosis and find its targets in host cells, we applied a genome wide screening of human targets for *Tg*ROP18 of RH strain with a Bi-molecular fluorescence complementation (BiFc) technique [25]. P2X1 was identified as a putative interacting partner of *Tg*ROP18. Our fluorescence resonance energy transfer (FRET) experiment further confirmed the interaction of *Tg*ROP18 with P2X1 in the cytoplasm (Figure 3A&B).

**Figure 3.**
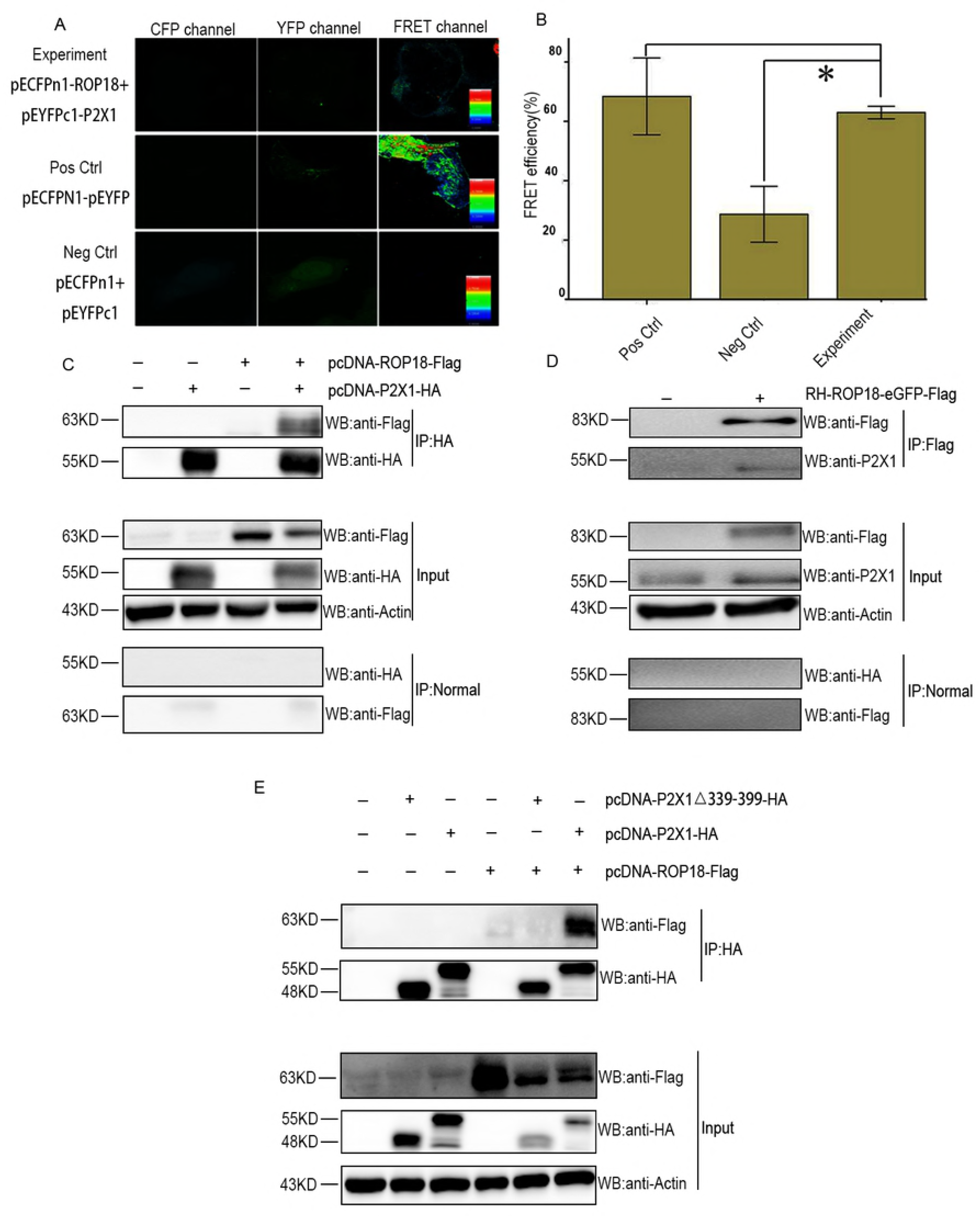
Identification of host cell P2X1 binding to *Tg*ROP18 by FRET and CO-IP. A: FRET identification of ROP18 binding to P2X1. COS7 cells were cultured in plates the day before transfection. pECFP-N1-ROP18^I^ and pEYFP-C1-P2X1 were co-transfected for the experimental group. pECFP-N1 and pEYFP-C1 were co-transfected for the negative control, and pEYFP-CFP was transfected for the positive control. The cells were stabilized for FRET experiment at 48 h post infection. The color matching the FRET signal intensity scale was shown in the FRET image of each transfection group. B: Quantitative analysis of FRET efficiency. The experimental group co-transfected with pECFP-N1-ROP18 and pEYFP-C1-P2X1 showed a significantly higher FRET efficiency than the negative control group. C: Lysates of COS7 cells transiently transfected with the indicated plasmids of pcDNA-ROP18-3×Flag and/or pcDNA-P2X1-HA were immunoprecipitated with the anti-HA antibody and detected by western blotting with the indicated antibodies. The result showed an association of ROP18 with P2X1 in both overexpressed cells. D: Lysates of SF268 cells infected with RH-ROP18-Flag(MOI= 3) for 36h or not were immunoprecipitated with anti-Flag antibody and detected by western blotting with the indicated antibodies. The result showed an association of endogenous ROP18 with P2X1. E: Lysates of COS7 cells transiently transfected with the indicated plasmids of pcDNA-ROP18-3×Flag and/or pcDNA-P2X1-HA, pcDNA-P2X1 Δ339-399- HA were immunoprecipitated with the anti-HA antibody and detected by western blotting with the indicated antibodies. The result showed that different form the full length P2X1, the mutant with P2X1-C terminus deletion did not bind with ROP18, which indicated that the intracellular carboxyl terminus of P2X1 was indispensible for its binding with ROP18.

To further assess the specificity of this interaction, we over-expressed flag-tagged *Tg*ROP18 together with HA-tagged P2X1 in COS-7 cells or infected SF268 cells with RH-ROP18-eGFP-Flag strain. The cell lysates were subjected to immunoprecipitation assay using an anti-HA antibody or anti-Flag antibody. The results indicated that not only the over-expressed Flag-tagged *Tg*ROP18 was immunoprecipitated by the overexpressed HA-tagged P2X1 (Figure 3C), but also the endogenous P2X1 from SF268 cells could be immunoprecipitated by endogenous *Tg*ROP18 tagged with Flag from RH-ROP18-eGFP-Flag strain (Figure 3D).

Previous studies have shown that the C terminus of P2X1 receptor plays an important role in the regulation of its expression and gating activity [17]. To understand whether this domain was involved in binding with *Tg*ROP18, we constructed a pcDNA-P2X1-HA plasmid with P2X1 C terminus (amino acids 339-399) deletion and co-transfected it with pcDNA-ROP18_Γ_3×Flag into COS-7 cells. The results showed that Flag-tagged *Tg*ROP18 could not be immunoprecipitated by HA-tagged P2X1 with C terminal deletion when compared with the HA-tagged full-length P2X1 which was used as a positive control (Figure 3E).

### 4-*Tg*ROP18 inhibited the P2X1-mediated apoptosis

To further identify if *Tg*ROP18 inhibited host cell apoptosis through binding with host cell protein P2X1, *Tg*ROP18 and/or P2X1 were over-expressed in COS7 cells for 48 hrs and stimulated with ATP for 24hrs. Cell apoptosis was then assessed with Annexin V/PI staining. Our results showed that over-expression of TgROP18 in COS7 cells significantly inhibited ATP-induced apoptosis (which was mediated by P2X1) when compared with the ATP treated cells (Figure 4A&C).

**Figure 4.**
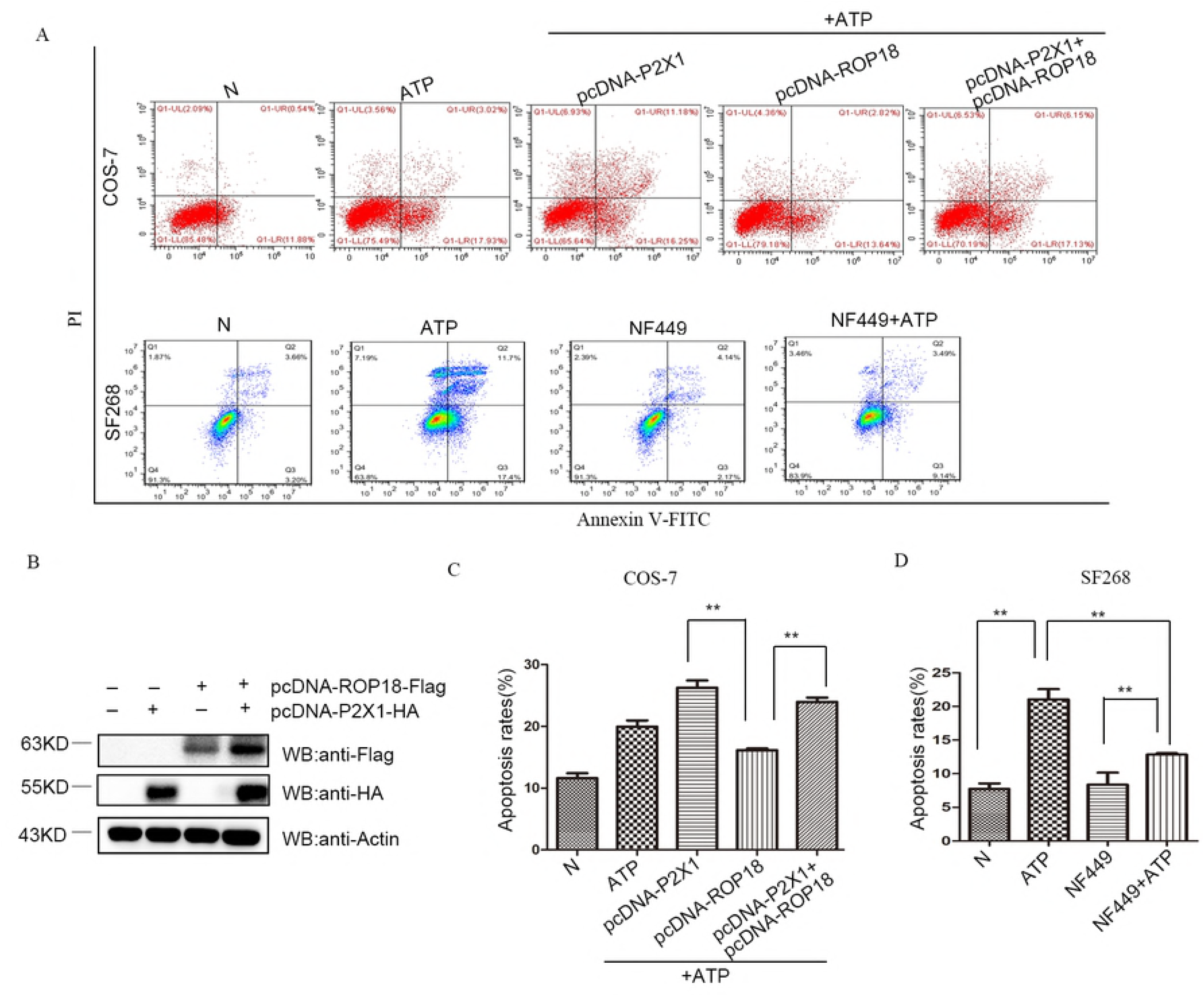
*Tg*ROP18 inhibited host cell apoptosis promoted by P2X1. Five groups of COS7 cells were transfected with pcDNA-ROP18 and/or pcDNA-P2X1, or not transfected, and then stimulated with ATP or not as indicated. Four groups of SF268 cells were treated with ATP and/or NF449 or not treated as indicated. A: The apoptosis rate of the cells in each group was detected by flow cytometry after Annexin V-FITC/PI staining. The results shown were the representative images of three independent experiments. B: The expression of Flag-tagged ROP18 and HA-tagged P2X1 was detected with western blot. **C&D:** The experiments were repeated for three times for one-way ANOVA statistical analysis, and the apoptosis rates of each group was expressed as mean ± SD. The result showed that P2X1 overexpression significantly promoted SF268 apoptosis, and *T. gondii* ROP18 overexpression significantly inhibited SF268 apoptosis induced by P2X1 overexpression (** P < 0.01) (Figure 4C). When the activity of P2X1 in ATP treated SF268 cells was inhibited with NF449, the cell apoptosis rate was significantly decreased compared to which of the ATP treatment group, and not significantly different from which of the normal control group (** P < 0.01) (Figure 4D).

When SF268 cells were pretreated with 4μM P2X1 specific inhibitor NF449 for 2hrs before ATP induction, the apoptosis rate in NF449+ATP treated group was significantly lower than which in ATP treated group, indicating that P2X1 was truly involved in the apoptosis induced by ATP, and functioned as a promoter for cell apoptosis (Figure 4D). On the other hand, we found the apoptosis index in NF449+ATP treated group was significantly higher than which in NF449 treated group, though both of the apoptosis indices in these two groups were significantly lower than which in ATP treated group. These results suggested that P2X1 inhibitor NF449 could not completely inhibit the apoptosis induced by ATP, and there probably existed some other proapoptotic proteins regulating on SF268 apoptosis induced by ATP besides P2X1.

### 5-*Tg*ROP18 inhibited host cell apoptosis through inhibition of P2X1-mediated Ca^2+^ influx but not through P2X1 degradation despite of their interaction

*T*gROP18 and P2X1 were over-expressed in COS7 cells for 72h, and the intracellular Ca^2+^ concentration was measured for 600s following addition of 60 μg/ml ATP into the culture medium. We found that P2X1 increased Ca^2+^ influx after ATP stimulation, and this process can be inhibited by over-expression of ROP18 of RH strain in the cells (Figure 5A&C). As a member of kinase family functioning in protein degradation, we next examined if ROP18 resulted in P2X1 degradation. We co-transfected pcDNA3.1(+)-ROP18-Flag of an increasing amount (0, 0.5, 1.0, 2.0 μg) with 2μg pcDNA3.1(+)-P2X1-HA into COS-7 cells for 48h. Our western blotting results showed that ROP18 expression did not affect the protein level of P2X1 (Figure 5C). All these evidences showed that ROP18 inhibited P2X1-mediated Ca^2+^ influx, as a result to inhibit cell apoptosis, but not through P2X1 degradation despite of their interaction.

**Figure 5.**
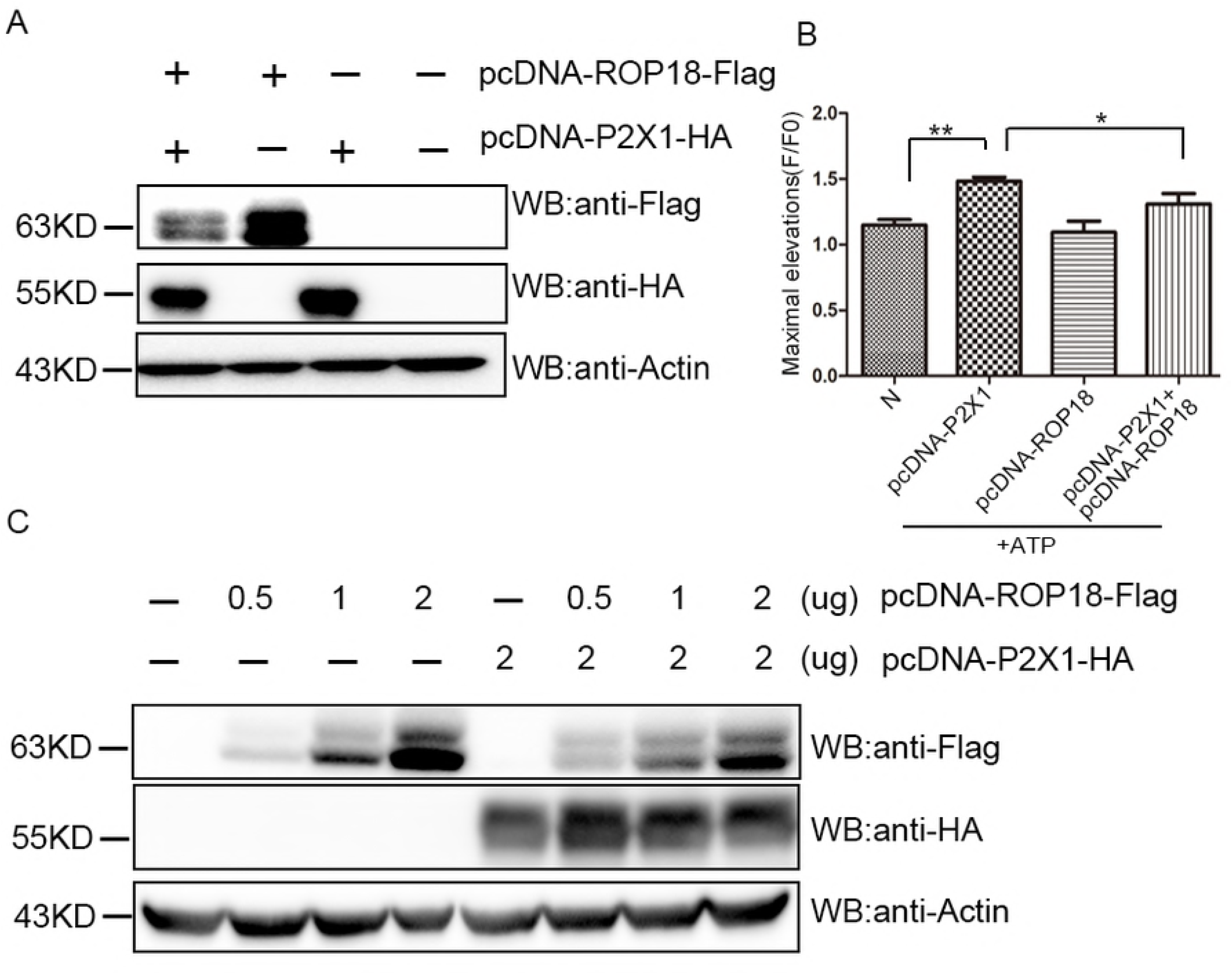
*Tg*ROP18 inhibited host cell apoptosis through inhibition of P2X1-medicated calcium influx but not through P2X1 degradation despite of their interaction. COS7 cells were transfected with 2.0μg of pcDNA-P2X1-HA and/or pcDNA-ROP18-3×Flag, or were transfected with a stable amount (2.0μg) of pcDNA-P2X1-HA and sequentially co-transfected with 0, 0.5, 1.0, and 2.0μg of pcDNA-ROP18-3×Flag respectively as indicated in each group. Ca^2+^ influx detection in each group were analyzed using flow cytometry after the cells were stimulated with ATP. The results shown were representative of three independent experiments. A: The expression of Flag-tagged ROP18 and HA-tagged P2X was detected with western blot. B: The experiments were repeated three times for one-way ANOVA statistical analysis, and the maximum elevation (F/F0) of the intracellular Ca^2+^ of each group was expressed as mean ± SD (* *P* < 0.05). The calcium influx level in P2X1 overexpressed cells was significantly higher than which in the cells with both P2X1 and ROP18 overexpression, which indicated that ROP18 significantly inhibited Ca^2+^ influx in SF268 cells. C: The result showed that ROP18 does not decrease P2X1 protein level, indicating that ROP18 didn’t degrade P2X1.

### 6-*Tg*ROP18 inhibited the depolarization of mitochondrion membrane and Cyt C translocation from mitochondria to cytoplasm in SF268 cells

Previous studies have shown that RH regulate the apoptosis of host cells mainly through mitochondrial pathway [26]. To identify whether this pathway was modulated by *Tg*ROP18 in *T. gondii* infection, we firstly detected the mitochondrial membrane depolarization after the SF268 cells were infected by RH or RH-Δ*ropl8* strains for 12hrs followed by ATP treatment for 12hrs. Both of our fluorescence microscope observation (Figure 6 A&B) and flow cytometry study (Figure 6 C&D) showed the intensity ratio of green fluorescence to red fluorescence was decreased in RH+ATP treated group when compared with ATP treated group, indicating that infection of RH strain inhibited mitochondrial membrane depolarization induced by ATP; Further, the intensity ratio of green fluorescence to red fluorescence was decreased in RH+ATP treatment group when compared with RH-*Δropl8* +ATP treatment group, implying that the depolarization of mitochondrion membrane was inhibited by *Tg*ROP18.

**Figure 6.**
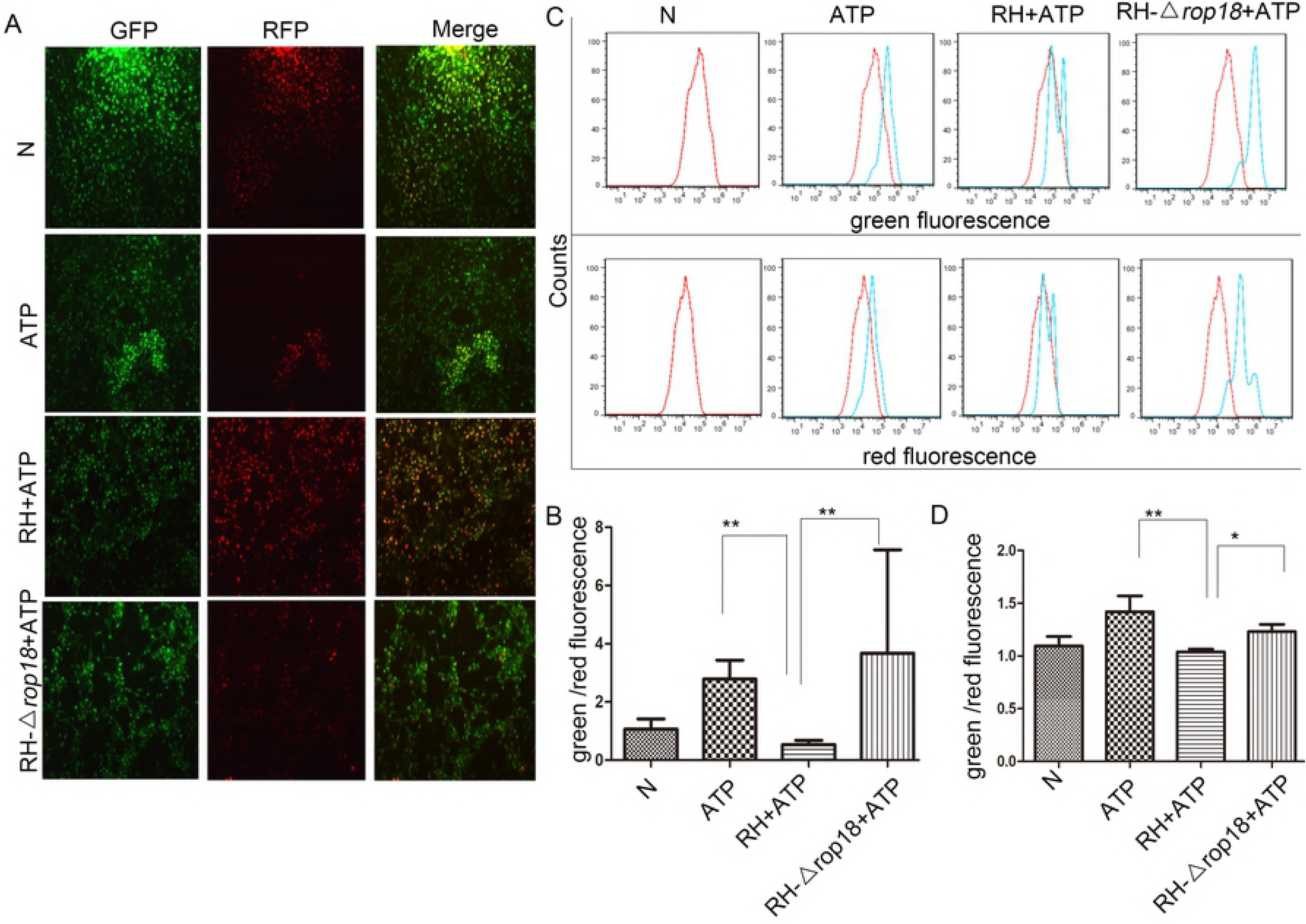
*T*⅛ROP18 inhibited ATP-induced mitochondrial depolarization in SF268 cells. The SF268 cells were infected with RH or RH-*Δropl8* tachyzoites at MOI of 13 or left uninfected, and then induced for apoptosis with 1 mg/ml ATP for additional 12h. A: The cells were stained with JC-1 and examined under a fluorescence microscope (10×) to determine the mitochondrial membrane depolarization. B: The cells were stained with JC-1 and examined under flow cytometry to determine the mitochondrial membrane depolarization. C: The percentages of green fluorescence to red fluorescence were determined by flow cytometry for each group of cells. Values are expressed as mean ± SD in each cell group, and the experiments were repeated three times for statistical analysis by one-way ANOVA (**P < 0.01, *P < 0.05). The result indicated that when treated with ATP, the RH-Δropl8 infected group showed significant mitochondrial depolarization compared to the RH infected group and the uninfected group.

Mitochondrial protein Cytochrome C (Cyt C) plays an important role in initiating the intrinsic apoptosis pathway. To evaluate the apoptotic result mediated by l· *Tg*ROP18, we next detected the situation of cytochromec (Cyt C) release from the mitochondria into the cytosol and the translocation of Bax (a proapoptotic Bcl2 family i protein) and Bcl2 (an antiapoptotic Bcl2 family protein) from the cytosol to the mitochondria in each group. In our western blotting results, relatively higher level of Cyt C was detected in the mitochondrial fraction, but lower level of Cyt C was detected in the cytosolic fraction in RH+ATP treatment group compared to which in I RH-Δ*rop/S*+ATP treatment group (P <0.01) (Figure 7 A, C1&C2). This result indicated that *Tg*ROP18 inhibited the cytochrome C translocation from mitochondria to cytoplasm.

**Figure 7.**
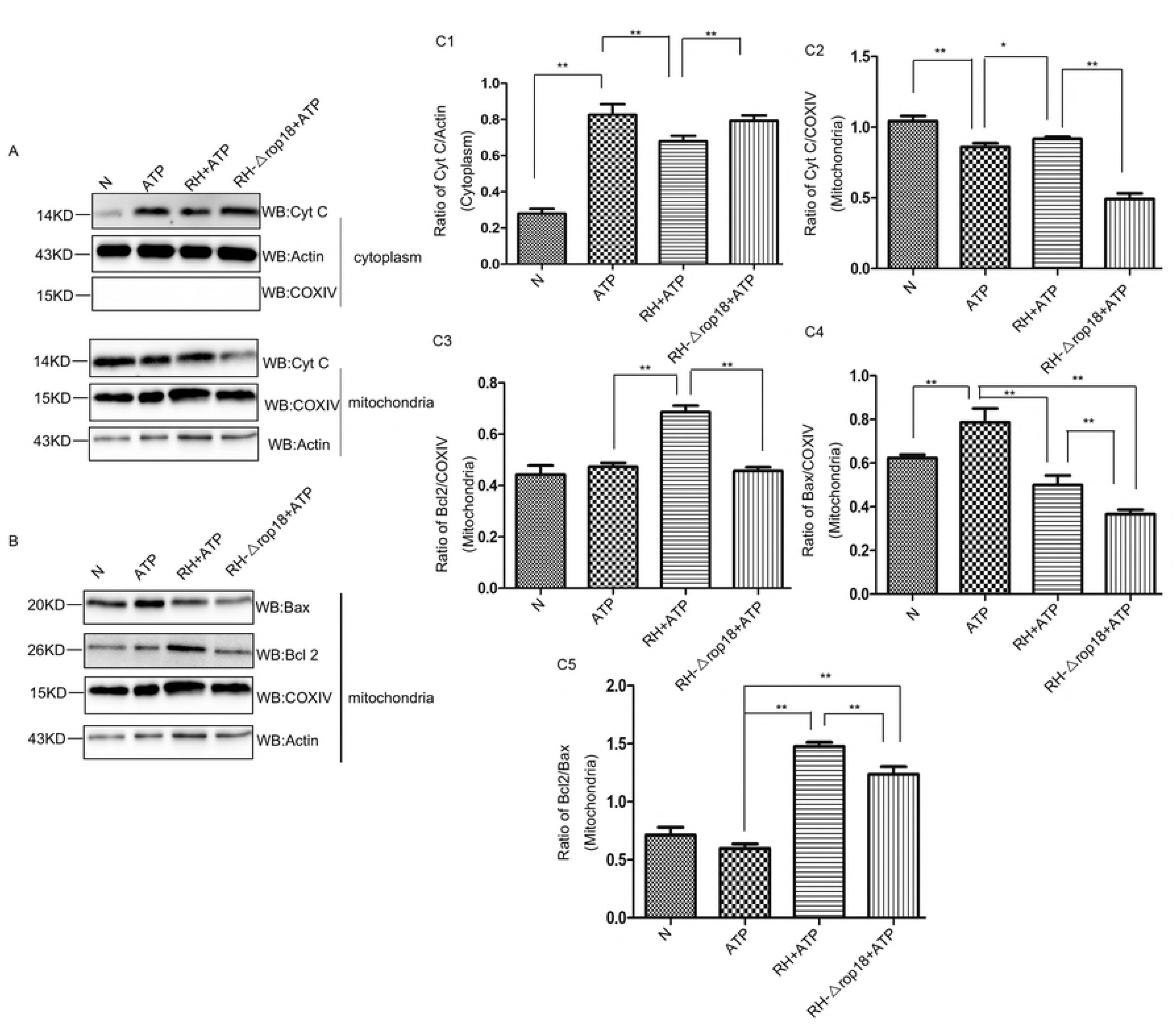
*T*gROP18 inhibits Cyt C release from mitochondria to cytoplasm in ATP treated SF268 cells. The SF268 cells were prepared as indicated in the legend of Figure 6. The cells were harvested and lysed, and the mitochondria and cytosol were fractionated. A&B: The mitochondrial and cytosol fractions were applied to western blot for detection of Cyt C, Actin and COXIV in cytoplasm and mitochondria fraction respectively, and detection of Bax, Bcl2 and COXIV in the mitochondria fraction. Data represented one of three times independent experiments. C: Densitometric analysis of western blotting using Image-J software was repeated three times. The ratios of Cyt C/actin, Cyt C/COXIV, Bcl2/COXIV, Bax/COXIV and Bcl2/Bax were calculated, and expressed as mean ± SD (** *P* < 0.01). The bar graphs showed that when induced by ATP in SF268 cells, RH infection significantly inhibited Cyt C translocation from mitochondria to cytoplasm (C1&C2), and increased Bcl2 and Bax’s translocation from cytoplasm to mitochodria (C3&C4) with higher Bcl2/Bax ratio in mitochondria fraction compared to RH-Δrop18 infection (C5). All these phenomena showed when SF268 cell apoptosis was induced with ATP, ROP18 significantly inhibited Cyt C translocation from mitochondria to cytoplasm and increased Bcl2/Bax ratio in mitochondria, and therefore inhibited the cell apoptosis.

Meanwhile, increased level of antiapoptotic protein Bcl2 and decreased level of proapoptotic protein Bax was found in the mitochondrial fraction of RH+ATP treated ¡ group compared to the ATP treated group (P<0.01) (Figure 7B, w), indicating that RH infection promoted the translocation of Bcl2 and inhibited the translocation of Bax from cytoplasm to mitochondria in SF268 cells. While the protein levels of mitochondrial Bax and Bcl2 were found both markedly higher in RH+ATP treated I group than which in RH-Δrop/S+ATP treated group (P<0.01) (Figure 7 B, C3&C4).

Further, we found that the Bcl2/Bax ratio in mitochondrial fraction was significantly higher in the RH+ATP group than which in the RH-Δrop/S+ATP group (P<0.01) ! (Figure 7C 5). These results suggested that though ROP18 promoted the translocation of both antiapoptoic protein Bcl2 and proapoptotic proteins Bax from cytoplasm to mitochondria, the ratio of Bcl2 to Bax in mitochondrial fraction was significantly ¡ increased by ROP18. As a result, the cell apoptosis was significantly inhibited by ROP18. Cytochrome C oxidase (COX) IV and actin were detected as the loading control for the mitochondrial fraction and the cytosolic fraction, respectively.

### 7-*Tg*ROP18 inhibited ATP-triggered caspases activation

The mitochondrial apoptotic pathway mostly depends on caspase-9, which in I turn activates the executioner caspases-3 and caspases-7 [27]. Our immunoblotting results indicated that in ATP treated SF268 cells, inactive full-length caspase-9 was cleaved into the active p35 and p37 fragments, inactive full-length caspase-7 was cleaved into active p20 fragments, and inactive full-length caspase-3 was split into p17 fragments. In SF268 cells treated with RH+ATP but not RH-Δ*ropl8*+ATP, significant inhibition of procaspase-9, procaspase-7 and procaspase-3 processing and activation to caspase-9, caspase-3 and caspase-7 was observed. Whereas no significant difference was detected in the levels of cleaved PARP (Poly ADP-ribose polymerase) between the ATP treated group and RH+ATP group (Figure 8).

**Figure 8.**
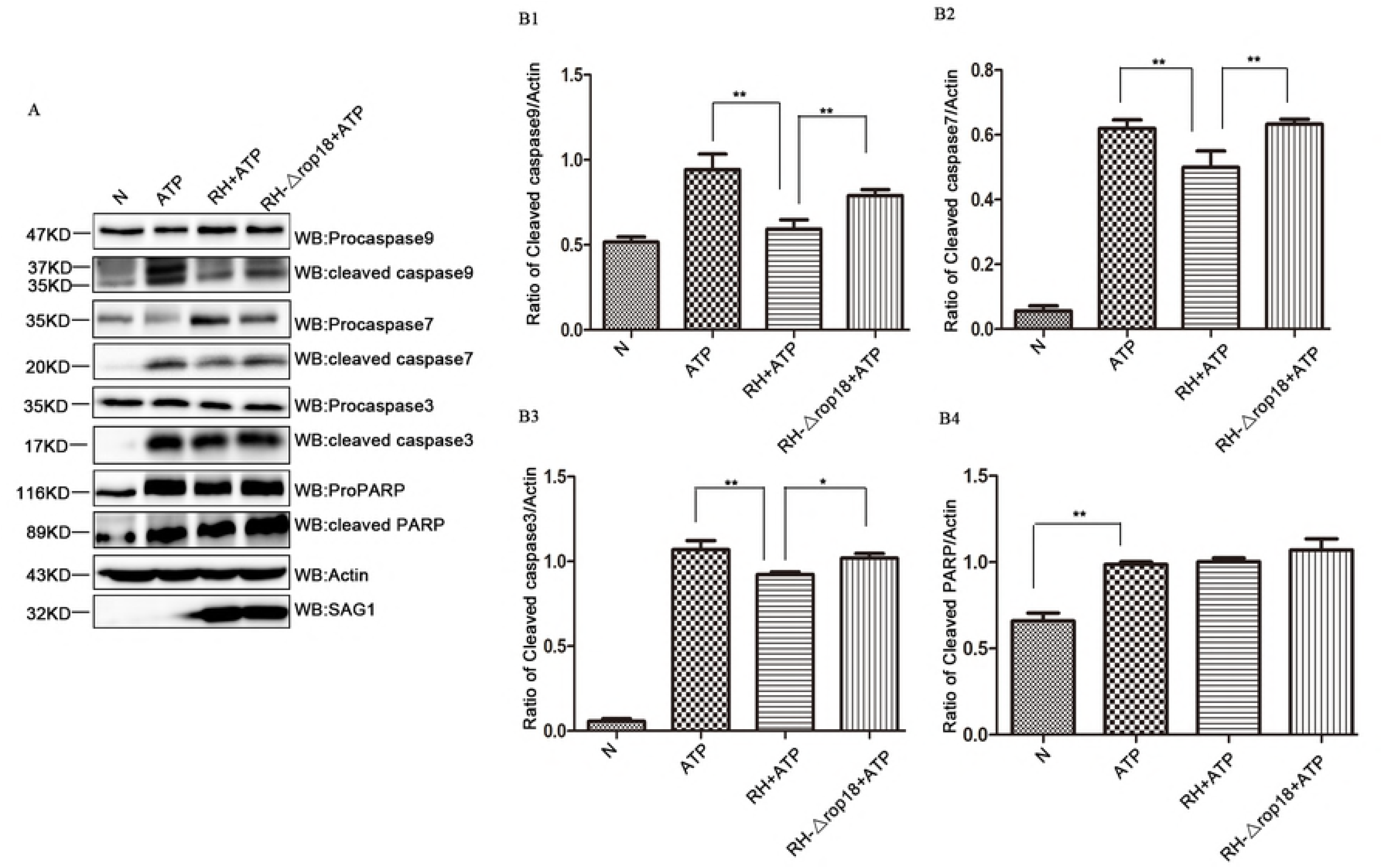
*Tg*ROP18 inhibited procaspase-3, procaspase-7 and procaspase-9 being cleaved to form caspase-3, caspase-7 and caspase-9 in ATP treated SF268 cells. The SF268 cells were prepared as indicated in the legend of Figure 6. The cells were harvested and lysed, the cell lysates were used for the western blot for detection of procaspase-3, procaspase-7, procaspase-9, pro-PARP, caspase-3, caspase-7, caspase-9 and PARP in SF268 cells. A: Antibody for caspase-9 recognized the full length 47 KD and cleaved form (35, 37 KD), antibody for caspase-7 recognized the full length 35 KD and the cleaved form 20 KD, antibody for caspase-3 recognized the full length 35 KD and the cleaved form 17 KD, and antibody for PARP recognized the full length 116 KD and cleaved form 89 KD. Actin was detected for loading control and SAG1 was detected for *T. gondii* tachyzoites amount, which showed consistent amount of host cells in each group and tachyzoites in each infection group. B: Densitometric analysis of western blot was conducted using Image J software, and the experiments were repeated three times for one-way ANOVA statistical analysis. The ratios of cleaved caspase-9 / actin, cleaved caspase-7 / actin, cleaved caspase-3 /actin, and cleaved PARP / actin were calculated and expressed as mean ± SD. (* *P* < 0.05). The graphs indicated that ROP18 significantly inhibited the cleavage of the full-length caspase-9, caspase-3 and caspase-7 to active forms, but had no effect on pro-PARP. All these caspases cleavage analysis showed that SF268 cell apoptosis induced by ATP was significantly inhibited by ROP18.

## Discussion

*Toxoplasma gondii* has a preference to infect the immune cells, hidden inside the immune cells, can it move across blood-brain barrier (BBB), then infects astrocytes and neurons [28]. Astrocytes can clear intracellular parasites through multiple mechanisms, while *T. gondii* can easily survive in neuro cells which lack full immune response capabilities [29]. Although it has been well accepted that *T. gondii* modulates host cell apoptosis during infection, it was still unclear whether this modulation was related to the strain virulence or not. Based on previously reported literatures, we found the apoptosis direction regulated by type I strain RH and type Π strain (NTE or ME49) in immune cells or neuro cells were varied (supplementary table S1). Therefore, we need to learn more about the host cell apoptosis resulted by infection of different types of *T. gondii.* We demonstrated in our research, when human glioblastoma cells (SF268), human monocyte/macrophage cells (THP-1) and murine macrophage cells (RAW264.7) were infected by RH, ME49 and VEG strains respectively for 6h or 28h, all of them showed a lower apoptosis rate compared to their uninfected controls, when cell apoptosis was induced by staurosporine (STS).

It has been reported that overexpression of *Tg*ROP18 significantly suppresses human embryo kidney epithelial cell 293T apoptosis induced by Actinomycin D at 48hrs post transfection [15], and *Tg*ROP18 targets P53 for degradation to inhibit host cell apoptosis [30]. On the contrary, *Tg*ROP18 is also reported to induce the apoptosis
of murine neuroblastoma N2a cells through the endoplasmic reticulum stress-mediated apoptosis pathway at 24hrs post infection [14]. In our research, we identified that *Tg*ROP18 significantly inhibited the apoptosis of SF268 cells induced by ATP when using RH and RH-Δ*rop18* strains in infection, but strangely this phenomenon was not seen in THP-1 and RAW264.7 cells. We further found that ROP18 interacted with human P2X1 which was endogenously expressed in SF268 cells but not in THP-1 and RAW264.7 cells (data were not shown). Probably it was the reason why ROP18 could inhibit the apoptosis induced by ATP through binding with P2X1 in SF268 cells, but not in THP-1 and RAW264.7 cells without P2X1 expression.

It has been known that RAW264.7 cell apoptosis can be induced by activation of protein kinase (ERK) and MAPK pathway regulated by extracellular signals through P2X4 as well as P2X7 receptors [31]; THP-1 cell apoptosis can be promoted via increase of intracellular calcium through P2X7 receptor [32]. Therefore, it seemed that ROP18 can only target at P2X1 but not P2X4 or P2X7 to regulate host cell apoptosis.

Owing to the insertion of a 2.1kb sequence in the promoter region of type III strains, ROP18 expression can nearly detected [33]. Whereas, we found in our study that type I strain VEG could also inhibited host cell apoptosis. It has been reported that ROP38, another member of rhoptry protein kinase (ROPK), is normally undetectable in virulent RH strain but is abundant in the avirulent VEG strain at the transcription level, and expression of ROP38 exerts a potent effect on the regulation of cell differentiation and apoptosis through mitogen-activated protein kinase (MAPK) pathway [34].

In our research, we identified that *Tg*ROP18 inhibited the apoptosis of SF268 cells through inhibition of mitochondria depolarization, translocation of Cyt C from mitochondria to cytoplasm, and ATP-triggered caspases activation. Cyt C is present in the mitochondrial inner membrane (MIM) media and plays a crucial role in transferring electrons [35]. Apoptotic signals such as DNA damage and nutrient deprivation increase the permeabilization of the MIM, and Cyt C will be released from the MIM media to the cytosol [36]. Cyt C in cytosol binds to apoptotic protease activating factor-1 (APaf-1) and forms a heptameric apoptosome complex to activate procaspase-9 to be cleaved to caspase-9, the activated caspase-9 stimulates the subsequent effecters caspase-3 and caspase-7 that eventually cause cell apoptosis [37,38]. Poly (ADP-ribose) polymerases (PARPs) are an important family of nucleoproteins which are mostly present in the nucleus and less in the cytosol. It is composed of three functional domains, namely central auto-modification domain, C-terminal catalytic domain and N-terminal DNA binding domain which assists in binding to both single-and double-stranded breaks in DNA for DNA repair [39,40]. In our study, pro-PARP can be activated into cleaved PARP when SF268 cells was treated by ATP for 12hrs. Activated PARP can repair damage DNA, that is why no DNA fragmentation was detected in in both infected or uninfected SF268 cells after induction with STS (data not shown)

The Bcl-2 family is classified into three groups according to their function in apoptosis and the number of Bcl-2 homology (BH) domains they possess. The anti-apoptotic Bcl-2 proteins and the proapoptotic proteins, Bax and Bak, all contain four BH domains BH1-BH4 [41]. In normal cells, Bcl-2 and Bax are located predominantly in the cytosol [42], but under apoptotic conditions, they will accumulate at the mitochondrial outer membrane [41,43]. Whether Cyt C is released into the cytosol partly depends on the ratio of Bcl2/Bax [44]. Our immunoblotting results showed that *Tg*ROP18 increased the ratio of Bcl2/Bax in the mitochondria, and then inhibited the translocation of Cyt C from the mitochondria to the cytoplasm.

In conclusion, this study identified that RH, ME49 and VEG infection inhibited the apoptosis of SF268, RAW264.7 and THP-1 cells when induced by STS. *Tg*ROP18 targeted C terminus of P2X1 and inhibit the apoptosis mediated by P2X1 in SF268 cells with P2X1 endogenous expression. When the cell apoptosis was induced by ATP, *Tg*ROP18 overexpression inhibited Ca^2+^ influx from the extracellular space to the cytoplasm in COS7 cells overexpressing P2X1. Furthermore, *Tg*ROP18 inhibited depolarization of mitochondrion membrane, Cyt C translocation from mitochondria to cytoplasm, and ATP-triggered caspase 9,7, and 3 activation in SF268 cells. All these findings supported that *Tg*ROP18 targeted host cell P2X1 and inhibited the cell apoptosis induced by ATP through the mitochondria pathway.

## Materials and methods

### Reagents

Apoptosis inducer: staurosporine (STS; S1421, Selleck, China), adenosine triphosphate (ATP) (A7699, Sigma-Aldrich, USA). Antagonist of P2X1: NF449(1391, TORICS, North America). Apoptosis detection: Annexin V-FITC apoptosis detection kit (KGA105-KGA108, KeyGen, China), JC-1 mitochondria membrane potential detection kit (KGA601-KGA604, KeyGen, China), Fluo-4 AM (Dojindo Laboratories, Japan). Cell Mitochondria Isolation Kit (C3601, Beyotime Biotechnology, China). Transfection reagents: Lipofectamine^®^ 3000 transfection kit (L3000015, Thermo Fisher Scientific, USA). Antibodies: Protein A-Agarose Immunoprecipitation Reagent (sc-2001) and normal rabbit IgG (sc-2028) were purchased from Santa Cruz Biotechnology; DDDDK-Tag Mouse mAb (AE005) was purchased from Abclonal. HA-Tag rabbit mAb (3724S), HA-Tag mouse mAb (2367), β-Actin rabbit mAb (13E5), caspase-3 rabbit mAb (9662), caspase-7 rabbit mAb (D2Q3L), caspase-9 rabbit mAb (9502), PARP rabbit mAb (9542) were all purchased from Cell Signaling Technology. Bax rabbit mAb (ab32503), Bcl2 rabbit mAb (ab32124), cytochrome C rabbit mAb (ab133504), P2X1 rabbit pAb (ab74053) and COXIV rabbit mAb (ab202554) were all obtained from Abcam.

### Cell culture

Human monocyte/macrophage cell line THP-1 (ATCC: American Type Culture Collection, USA) and Human glioblastoma cell line SF268 (ATCC) were propagated in RPMI 1640 (Thermo Fisher Scientific) supplemented with 10% (v/v) fetal bovine serum (Gibco, Australia), 100 U/mL penicillin and100 μg/mL streptomycin at 37 °C and 5% CO_2_. Mouse monocyte/macrophag cell line RAW264.7 (ATCC),
*Cercopithecus aethiops* kidney fibroblast cell line COS7 (ATCC), and human foreskin fibroblast cell line HFF(ATCC) were all cultured in DMEM supplemented with 10% (v/v) fetal bovine serum (Gibco, Australia), 100U/mL penicillin and100 μg/mL streptomycin at 37 °C and 5% CO_2_.

### *T. gondii* culture, cell infection and induction of apoptosis

Type I strain RH, RH-*Δrop18,* RH-ROP18-eGFP-Flag, type II strain ME49 and type III strain VEG were cultured in HFFs in DMEM (Thermo Fisher Scientific) supplemented with 1% (v/v) FBS (Gibco, Australia), 100 U/ml penicillin, and 100μg/ml streptomycin (Invitrogen) at 37 °C and 5% CO2. At 3-5 days, when most of the cells were ready to be ruptured by the tachyzoites. The cells were scraped and harvested to pass through a syringe for multiple times. The tachyzoites were then purified by passing through a 3-μm filter (Whatman, England), then pelleted at 2500g for 10min and resuspended in DMEM medium. The tachyzoites were counted with hemocytometer.

Before infection, THP-1 were cultured in the presence of 100 ng/ml phorbol 12-myristate 13-acetate (PMA) (ab120297, Abcam, England) for 24 h for differentiation into macrophages as described previously [45]. THP-1, RAW264.7 and SF268 cells were infected with RH, ME49 and VEG at multiplicity of infection (MOI) of 3 and incubated for 2 h or 22 h at 37°C in 5% CO_2_. Before apoptosis induction, cells were washed with PBS to remove non-adherent parasites, then, THP-1 cells were treated with 2μM STS, SF268 cells were treated with 350nM STS and RAW264.7 cells were treated with 250nM STS for 4 h or 6 h, Then, The apoptotic level of the cells was analyzed with AnnexinV-FITC/PI and DNA fragmentation.

RH-wild type and RH-Δ*rop18* tachyzoites were separated from the host cells by centrifugation, and SF268, THP-1, RAW264.7 cells were infected with the tachyzoites (MOI =13). The cells were washed with PBS at 12 h post infection. and then the cells were treated with 1mg/ml adenosine triphosphate (ATP) for an additional 12 h to induce apoptosis. The apoptotic level of the cells was analyzed with AnnexinV-FITC/PI.

### Flow cytometry detection of apoptosis with Annexin V-FITC/PI

The apoptotic levels of RAW264.7, THP-1 and SF268 cells were determined with isothiocyanate (FITC)-Annexin V/propidium iodide (PI) kit (KeyGen, China). Cells were briefly digested with 0.25% trypsin (Thermo Fisher Scientific, USA) and washed twice with cold PBS, then resuspended in 500 μL binding buffer, 5 μL FITC-conjugated Annexin V and 5 μl propidium iodide (PI) were added to the suspension and incubated at room temperature for 15 min in the dark. The flow cytometer (BD FACSCalibur, USA) was used for data collection, FlowJo software was used for data analysis. According to the kit manual, Annexin V-FITC+/PΓ cells represent the early apoptotic cells, and AnnexinV-FITC+/PI+ cells are the late apoptotic cells [46,47].

### DNA fragmentation detection

The extraction of DNA fragments followed the previous reported processes [48]. Cells were briefly digested with 0.25% trypsin and washed twice with pre-cold PBS. DMSO was added to the cell pellet and immediately vortexed, then, an equal volume of TE buffer (pH7.4) with 2% SDS was added to the cell suspension followed by vortexing. The solution was centrifuged at 12000 *g* at 4°C for 10 min and 30μl supernatant was loaded on agarose gels for electrophoresis.

### Fluorescence resonance energy transfer (FRET) experiment

COS7 cells were cultured in plates the day before transfection. pECFP-N1-ROP18¡ and pEYFP-C1-P2X1 were co-transfected for the experimental group. pECFP-N1 and pEYFP-C1 were co-transfected as a negative control and pEYFP-CFP was transfected as a positive control [49]. Forty-eight hours post transfection, the cells were fixed in 4% paraformaldehyde at 37°C for 30 minutes and then washed twice with PBS. The FRET efficiency of different transfection groups were measured under a confocal laser scanning microscope (Olympus FLUOVIEW FV1000) [50].

### CO-Immunoprecipitation analysis

SF268 and COS-7 cells were grown in T75 flasks to 100% confluence. SF268
cells were infected with RH-ROP18-eGFP-Flag strain or not for 36 h. COS-7 cells were transfected with pcDNA3.1(+)-ROP18-3×Flag and/or pcDNA3.1(+)-P2X1-HA or with pcDNA3.1(+)-ROP18-3×Flag and/or pcDNA3.1(+)-P2X1Δ339-399-HA for 48h. The cells were washed with PBS for 3 times and lysed using cell lysis buffer (P0013, Beyotime, Shanghai, China) with 1 mM phenylmethanesulfonyl fluoride (WB-0181, Beijing Dingguo Changsheng Biotechnology, Beijing, China). Cell lysates were centrifuged at 14,000 *g* for 10 min at 4 °C. The supernatants were harvested and incubated with the primary antibody anti-HA antibody (3724S, CST), Monoclonal anti-FLAG antibody (F1804, Sigma Aldrich) or rabbit normal IgG for 2h at 4 °C with gentle rotation. Then Protein A-Agarose (Santa Cruz) was added to the mix and incubated overnight at 4°C with gentle rotation. Beads were collected by centrifugation at 500 *g* for 5 min at 4°C, and then resuspended in SDS-PAGE sample buffer. The samples were boiled and loaded to the gels for SDS-PAGE, and then analyzed by WB [51].

### ROP18 inhibited host cell apoptosis mediated by P2X1

COS-7 cells were transfected with pcDNA3.1(+)-ROP18-3×Flag and/or pcDNA3.1(+)-P2X1-HA. At 48 hours PI, the cells were treated with 60μg/ml ATP for 24h to induce cell apoptosis or not induced for control. SF268 cells were treated with 4μM NF449 (P2X1 receptor antagonist) for 2hrs or not treated, then subjected to 1mg/ml ATP treatment for 12hrs. The normal control cells were treated with neither NF449 nor ATP. The apoptosis rate of the cells was analyzed with AnnexinV-FITC/PI.

### Detection of Calcium influx with Fluo-4 AM

COS7 cells were grown in 24-well plates and co-transfected with pcDNA3.1(+)-ROP18-Flag and/or pcDNA3.1(+)-P2X1-HA plasmids for 72h. The cells were then harvested and washed twice with Hank’s Balanced Salt Solution (HBSS) (Invitrogen). Two hundred and fifty μL of 5 μM Fluo-4 AM (Dojindo Laboratories, Japan) (an indicator of intracellular calcium ions) was added to the cells and incubated for 30 min at 37 °C. After being washed twice with HBSS, the cells were analyzed with flow cytometry. Fluo-4 AM is virtually non-fluorescent, acetoxymethyl (AM) ester moiety allows this dye to cross the cell membrane, whereupon endogenous esterases cleave the AM group to liberate the active dye and Ca^2+^ can bind with Fluo-4 intracellularly to emit green fluorescence. Time series reads were taken every 2 seconds for 100s and the mean value was calculated as the baseline read (F0). Following 60μg/ml ATP stimulation, the Green fluorescence intensity reflecting the free intracellular calcium ions level in each group was recorded for 500s with flow cytometry. The intracellular calcium level was quantified by the ratio of strongest signal fluorescence signal to the basal signal (F/F0).

### Detection of mitochondrial membrane depolarization with JC-1 staining

SF268 cells were infected with RH or ΔROP18-RH tachyzoites (MOI=13). Twelve hours later, 1 mg/ml ATP was added to the cells to induce apoptosis for 12 h. Cells were then digested with 0.25% trypsin and rinsed with PBS twice. Five hundred micro liters of incubation buffer containing 1 μL of JC-1 was added to the cells and incubated for 15 min at 37°C in 5% CO2. The cells were pelleted and washed with 1×incubation buffer twice, the cells then were visualized under a fluorescence microscope (Nikon, Japan) and flow cytometry (BD Biosciences, USA). For fluorescence microscope with excitation at 550 nm and emission at 600 nm for red fluorescence and excitation at 485 nm and emission at 535 nm for green fluorescence. For flow cytometry, JC-1 was excited with the 488 nm argon laser, and JC-1 green and red fluorescence were recorded on FL1 and FL2 channels. A minimum of 10000 cells within the gated region were analyzed. The value was calculated by the ratio of green fluorescence to red fluorescence. The lipophilic, cationic dye JC-1 staining can discriminate polarized and depolarized mitochondria because in polarized mitochondria the normal green fluorescent JC-1 dye forms red fluorescent aggregates in response to their higher membrane potential, while in depolarized mitochondria, the red fluorescent aggregates convert to monomeric form and exhibit green fluorescence [52].

### Separation of cytosolic proteins and mitochondrial protein in SF268 cells

The isolation of mitochondrial protein and cytosol proteins were performed using the Cell Mitochondria Isolation Kit. SF268 cells were harvested and washed twice with cold PBS, and then incubated in 100 μL ice-cold mitochondrial lysis buffer with gentle rotation at 4 °C for 15 min. Cell suspension was then transferred into a glass homogenizer and homogenized for 30 strokes using a tight pestle on ice. The homogenate was subjected to centrifuging at 600 *g* for 10 min at 4 °C to remove nuclei and unbroken cells. Then, the supernatant was collected and centrifuged again at 12,000 *g* for 10 min at 4 °C to obtain the cytosolic (supernatant) and mitochondrial (deposition) fraction. Proteins of Cyt C, Bcl2 and Bax in the cytosol and mitochondria were detected by western blotting. The intensity of Cyt C, Bcl2 and Bax bands with non-saturated exposure from three independent experiments were analyzed using Image J software, and the proportion of Cyt C, Bcl2 and Bax to the loading control Actin or COXIV were calculated.

### Western blotting

Protein samples were diluted with 6× SDS PAGE loading buffer and then boiled for 10 min. Boiled samples were loaded to 15% SDS-polyacrylamide gels for electrophoresis, and then transferred to polyvinylidene fluoride (PVDF) membrane. After transferring, the PVDF membrane was blocked in PBS/5% BSA/0.05% Tween-20 at 37 °C for 2 h with gentle shaking, and then probed with the primary antibody under the same condition as blocking. Then, the membrane was incubated with the secondary antibody. The PVDF membranes were visualized by enhanced chemiluminescence (ECL) detection as recommended by the manufacturer.

### Statistical analysis

All experiment were repeated for three times and data were collected for statistical analysis. Data were analyzed using GraphPad Prism 5 (San Diego, CA) and presented as mean ± SD. Statistical significance was determined using a one-way ANOVA test, and *\*P*<0.05 and *\*\*P*<0.01 was considered as significant difference.

## Acknowledgements

The authors are grateful to the participants in this study and the anonymous reviewers and editors for their comments and valuable inputs.

---

**Figure S1. Infection with *T. gondii* RH, ME49 and VEG inhibits STS-induced apoptosis of RAW264.7, THP-1 at 6h post infection.**
The indicated cells were infected with RH, ME49 and VEG strains, respectively, at MOI (multiplicity of infection) of 3 for 2h and treated with staurosporine (STS) for 4 h. The cells were collected for apoptosis analysis by flow cytometry (FCM) and DNA fragmentation. A: Apoptotic cells included the early apoptotic cells (Annexin V^+^/ PI^−^) and the late apoptotic cells (Annexin V^+^/ PI^+^). B: The quantitative data converted from FCM data. Values were expressed as mean ± SD in each group, and the experiments were repeated three times for one-way ANOVA statistical analysis (**P < 0.01). C: DNA ladder assay of the indicated cells (RAW264.7 and THP-1). 30μl of lysate from each group was loaded on 1.5% agarose gel for electrophoresis. M: 100bp DNA ladder. Both the FCM data and the DNA fragmentation detection showed the apoptosis of RAW264.7 and THP-1 cells induced by STS were significantly blocked by infection of RH, ME49 and VEG, respectively for 6h.

**Figure S2. Infection of *T. gondii* RH, ME49 and VEG inhibits the STS-induced apoptosis of RAW264.7 and THP-1 cells at 28h post infection.**
The indicated cells were infected with RH, ME49 and VEG strains, respectively, at MOI of 3 for 22h and treated with STS for 6h. The cells were collected for apoptosis analysis by flow cytometry (FCM) and DNA fragmentation. A: Apoptotic cells included the early apoptotic cells (Annexin V+/ PI^−^) and the late apoptotic cells (Annexin V^+^/ PI^+^). B: The quantitative data converted from FCM data. Values were expressed as mean ± SD in each group, and the experiments were repeated three times for one-way ANOVA statistical analysis (**P < 0.01). C: DNA ladder assay of the indicated cells (RAW264.7 and THP-1). 30μl of lysate from each group was loaded on 1.5% agarose gel for electrophoresis. M: 100bp DNA ladder. Both the FCM data and the DNA fragmentation detection showed the apoptosis of RAW264.7 and THP-1 cells induced by STS were significantly blocked by infection of RH, ME49 and VEG, respectively for 28h.

**Figure S3.*T. gondii* major virulence factor ROP18 didn’t inhibit RAW264.7 and THP-1 cell apoptosis induced by ATP.**
The RAW264.7 and THP-1 cells were infected with RH or RH-*Δropl8* tachyzoites (MOI=13) or left uninfected for normal control (N) or positive control (ATP treatment). At 12 h post infection, 1 mg/ml ATP was added to the cells for additional 12 h except the normal control group. A: The apoptosis of the SF268 cells from each group was detected by flow cytometry after Annexin V-FITC/PI staining. B: The percentages of apoptotic cells were separately determined for each group of cells. Values were expressed as mean ± SD in each cell group, and the experiments were repeated three times for one-way ANOVA statistical analysis (**P < 0.01). The RAW264.7 and THP-1cells infected by RH strain showed no significant different apoptosis rate compared to the RH-*Δropl8* infection group, which implied that ROP18 had no effect on the cell apoptosis induced by ATP in RAW264.7 and THP-1 cells.

**Tabel S1 The summarized information of *Toxoplasma gondii* modulating immune and neuro cell apoptosis**

